# Zinc and Cobalt exposure influence *Akkermansia muciniphila* growth and short-chain fatty acid metabolism

**DOI:** 10.1101/2025.08.06.668965

**Authors:** Shoshannah Eggers, Joseph Eggers, Maria J. Banas, Chris Gennings, Vishal Midya

**Author notes:** Corresponding Author: Shoshannah Eggers.

## Abstract

*Akkermansia muciniphila* is a probiotic bacterium that has been proposed as a potential intervention for depression. *A. muciniphila* is a common mucin degrader, producing short fatty chain acids (SCFAs), including acetate, butyrate, and propionate, which can help regulate mood via the gut-brain axis. However, the interplay between *A. muciniphila* and other neuroactive environmental exposures is unclear. The objective of this study was to assess the effects of Zn and Co exposure, individually and together, on *A. muciniphila* growth and SCFA production. Minimum Inhibitory Concentration (MIC) was calculated for each metal, and sub-MIC exposure testing at high and low levels was conducted with single and combined metals. Growth curves were measured for each exposure condition, and SCFAs were measured using liquid chromatography-mass spectroscopy. Growth and SCFA production were compared between exposures and controls using paired t-tests and linear regression models. All p-values were corrected using FDR adjustments (q-values). *A. muciniphila* was tolerant to both metal exposures, with MICs at 750mg/L for Zn and 4,000 mg/L for Co. Growth at 24-and 48-hours was only significantly reduced upon exposure to high levels of Co, alone and in combination with Zn. Production of propionate significantly increased with high and low Zn exposures alone and in combination with low Co exposure (q-values <0.01), and significantly decreased with high Co exposure alone and in combination with Zn (q-values <0.01). Overall, *A. muciniphila* was metal tolerant, supporting its use as a probiotic in the presence of other neuroactive metal exposures.

## 1. INTRODUCTION

Mental health disorders affect 1 in 5 US adults and arise from complex and heterogeneous etiologies.(National Institute of Mental Health, 2024) Both metal exposures and the gut microbiome have been widely shown to contribute to mental health outcomes,(Li et al., 2018; Liu et al., 2023; Sun et al., 2023) however, the way in which metals and microbes interact in this context is unclear. Probiotic gut microbes have been explored as a potential intervention for depression and anxiety working through the gut brain axis, with mixed results.(Dinan and Cryan, 2017; Chen et al., 2024) Several reviews have made a point of the difficulty in assess the efficacy of probiotics interventions for depression given the variety of probiotic organisms used, dose and duration of treatment, and target population. (Cohen Kadosh et al., 2021; Hofmeister et al., 2021; Ng et al., 2018) This high level of variability suggests a need for a more targeted probiotic approach. Given the influence of metal exposures on depression, a precision environmental health approach could be used to identify subsets of the population that are highly susceptible to depression based on their exposure patterns, which could be paired with a probiotic bacteria treatment that would be most effective as an intervention for those subsets. However, greater understanding of the interactions between those metals and probiotic microbes is needed to develop these precision interventions.

*Akkermansia muciniphila* is a probiotic bacteria that has shown promising results as an intervention for depressive symptoms in animal models.(Cheng et al., 2022; Ding et al., 2021; Guo et al., 2022, 2024; Sun et al., 2023) *A. muciniphila* is among the top 20 bacterial species that inhabit the human gut throughout the lifespan.(Collado et al., 2007; Derrien et al., 2008) *A. muciniphila* symbiotically colonizes the mucosal layer of the intestinal epithelium and breaks down mucins, which are high-molecular-weight glycoproteins.(Allen, 1983; Liu et al., 2022) This mucin degradation leads to *A. muciniphila’s* production of short-chain fatty acids (SCFAs) via anaerobic fermentation.(Derrien et al., 2008; Liu et al., 2022) SCFAs, including acetate, butyrate, and propionate, are exclusively produced by bacteria, and serve a range of functions in the human body, including neuromodulation.(Morrison and Preston, 2016) SCFAs modulate the release and function of neurotransmitters in the brain, including serotonin,(Mirzaei et al., 2021; Ruhé et al., 2007) which plays a crucial role in regulating mood.(Baldwin and Rudge, 1995) SCFAs also have systemic anti-inflammatory properties.(Vinolo et al., 2011) Chronic inflammation has been associated with an increased risk of mood disorders, including anxiety and depression,(Li et al., 2019) and SCFAs can help maintain a balanced immune response, reducing the risk of excessive inflammation that may contribute to depressive symptoms.(Hao et al., 2019) SCFAs also help maintain blood-brain-barrier integrity by inducing the production of tight junction proteins that strengthen the barrier between epithelial cells, protecting the brain from harmful substances and inflammatory molecules. (Fock and Parnova, 2023; Tang et al., 2022) Little is known about the effects of metal exposure on *A. muciniphila* and it’s ability to produce SCFAs. However, metal exposures are consistently associated with altered composition and function of the gut microbiome as a whole.

A growing body of evidence suggests that metal exposures have negative impacts on the human gut microbiome, including alterations in SCFA synthesis. In animal models, metal exposures alter the abundance of specific bacterial taxa, gut microbial diversity, and metabolic profiles, such as reduced SCFA synthesis.(Gao et al., 2017; Sauer and Grabrucker, 2019; Zhang et al., 2017) Our group has demonstrated associations between metal exposures and the human gut microbiome using epidemiologic data from multiple cohorts,(Eggers et al., 2023, 2019; Sitarik et al., 2020) and further linked metals and gut microbes to inflammation and depressive symptoms in late childhood.(Midya et al., 2024c, 2024a) Using a novel analytical method called Microbial Co-occurrence Analysis (MiCA)(Midya et al., 2023) we identified a negative association between a signature of prenatal metal exposure (high Zn and low Cr in the second trimester, and low Co in the third trimester) and depression in childhood, that is moderated by the presence of *A. muciniphila* in the gut microbiome in childhood.(Midya et al., 2024b) However, to better understand the underlying biological mechanisms of this association and moderation, more direct biochemical information about the interaction between these metals and *A. muciniphila* is needed. The purpose of this study was to establish a baseline understanding of the relationship between metal exposures and the growth and SCFA production of *A. muciniphila* to inform its potential effectiveness as a probiotic in the presence of neuroactive metal exposures. In this study, we measured the growth and SCFA production of *A. muciniphila* in the presence of Zn and Co individually and as a mixture.

## 2. METHODS

### 2.1. Establishing *A. muciniphila* cultures

Certified starter cultures of *A. muciniphila* were purchased from ATCC (Manassas, VA).

All culture and enrichment work was conducted in an anaerobic chamber (Bactron 300, Sheldon Manufacturing, Cornelius, OR) as *A. muciniphila* is a strict anaerobe, and SCFAs are products of anaerobic fermentation. A modified Brain Heart Infusion (BHI) enrichment broth with added cysteine, yeast, and mucin was used for all cultures. An optimal growth curve was determined by repeated optical density (OD) measurement, and strain identity was confirmed using PCR with *A. muciniphila* 16S rDNA primers 5′-CAGCACGTGAAGGTGGGGAC-3′ and 5′-CCTTGCGGTTGGCTTCAGAT-3′.(Kim et al., 2021)

Pilot growth curves were measured every 6 hours for 72 hours to determine timing of the lag, exponential, stationary, and decline phases of the growth curve occurred. Based on those results, growth measurements under experimental conditions were taken at 0 hours (baseline lag phase), 12 hours (beginning of exponential phase), 24 hours (peak of exponential phase), and 48 hours (stationary phase). For the SCFA measurement, three 300uL aliquots were taken at each time point, for every growth condition. Aliquots were flash frozen on liquid nitrogen and stored at -80oC.

### 2.2 Metal Exposure

Zinc citrate was used for Zn exposure (National Center for Biotechnology Information, 2023; Wegmüller et al., 2014) and cobalt (II) chloride was used for Co exposure (Finley et al., 2013) because both are bioavailable, soluble, and common forms of human exposure.

We first determined the minimum inhibitory concentration (MIC) of each metal individually using a series of titer plates with increasing metal concentrations mixed into the enrichment broth.(Eggers et al., 2021; Kafilzadeh et al., 2012) MICs were considered the minimum concentration at which there was no bacterial growth in the titer plate as detected by the unaided eye or OD reading.(White et al., 1996) We conducted MIC testing in two replicates for each metal. Using the MIC for each metal as a guide, we chose a high and low concentration for both Zn and Co, all of which were at a level with altered growth of *A. muciniphila*, but lower than the MIC. For co-exposure, we used the same two low and high concentrations for each metal in all possible combinations (**Table 1**). Each exposure condition was grown in triplicate with one negative control (metal in the media with no bacteria inoculation) and one positive control (bacteria inoculation with no metal in the media) also in triplicate.

**Table 1.** Description of each exposure test condition and controls.

| Label | *A. muciniphila* | Co Level | Zn Level |
| --- | --- | --- | --- |
| Positive Control | Yes | None | None |
| Total Negative Control | No | None | None |
| Zn- Test | Yes | None | Low |
| Zn- Negative Control | No | None | Low |
| Zn+ Test | Yes | None | High |
| Zn+ Negative Control | No | None | High |
| Co- Test | Yes | Low | None |
| Co - Negative Control | No | Low | None |
| Co + Test | Yes | High | None |
| Co + Negative Control | No | High | None |
| Zn-/Co- Test | Yes | Low | Low |
| Zn-/Co- Negative Control | No | Low | Low |
| Zn-/Co+ Test | Yes | High | Low |
| Zn-/Co+ Negative Control | No | High | Low |
| Zn+/Co- Test | Yes | Low | High |
| Zn+/Co- Negative Control | No | Low | High |
| Zn+/Co+ Test | Yes | High | High |
| Zn+/Co+ Negative Control | No | High | High |

Tests and negative controls include metal at these concentrations: Zn- (Low) is 200mg/L; Zn+ (High) is 500mg/L; Co- (Low) is 200mg/L; Co+ (High) is 3,500mg/L. Test conditions included like *Akkermansia muciniphila* starter cultures, and negative controls did not contain any bacteria. Negative control Labels with two metals included both metals at their respective concentrations indicated by the + or – symbols.

### 2.3. SCFA Measurement

Flash frozen aliquots from 12- and 24-hours growth were sent to the Metabolomics Core Facility at the University of Iowa Carver College of Medicine. Samples were processed using 4-fold (weight/volume) extraction solvent (Acetonitrile:Methanol:Water (2:2:1)) containing deuterated SCFA standards. Samples were rotated at 20 oC for 1 hour and then centrifuged at 21,000 g for 10 min. The supernatant was used for LC-MS analysis using a Thermo Q Exactive hybrid quadrupole Orbitrap mass spectrometer with a Vanquish Flex UHPLC. 3 uL of sample was injected into a ZIC-pHILIC LC column (50 × 2.1 mm). The mass spectrometer was operated in negative full-scan mode from 0.05 to 4.55 minutes. Acquired LC-MS data were processed by Thermo Scientific TraceFinder 4.1 software, and metabolites were identified based on the facility standard-confirmed, inhouse library.

Analyte signal was corrected by normalizing to the deuterated analyte signal and the signal obtained from the negative control.

### 2.4. Statistical Analysis

All statistical analysis was completed using R. The packages used for data manipulation, visualization, and analysis included ggplot3, tidyr, readr, dplyr, stringr, and broom. (Robinson et al., 2025; Wickham et al., 2025, 2024b, 2024a, 2023a, 2023b) All unadjusted p-values were FDR corrected using q-values,(Storey et al., 2004) with a q-value of 0.05 indicating statistical significance.

#### 2.4.1. *A. muciniphila* Growth Curve Analysis

Growth curve data was computed with estimated mean growth and standard error (SE) over time. For visualization, line plots were created to display the growth of Akkermansia over time, stratified by exposure conditions and with computed means and SE. Growth patterns of exposure test conditions and controls were analyzed and mapped at 0, 12, 24, and 48 hours to capture the beginning of the lag phase, beginning of the exponential phase, peak of the exponential phase, and stationary phase, respectively. The mean (SE) of growth differences was computed by subtracting the positive control values from the reference exposure values. Given the replicates, Paired t-tests were used to ascertain any time-point-wise statistical significance. The growth rate was estimated using simple linear regression models.

#### 2.4.2. *A. muciniphila* SCFA Analysis

Summary statistics were grouped by exposure condition, metabolite, and growth period. Mean metabolite concentrations and SE were calculated. Propionate, acetate, and butyrate were analyzed for each exposure condition at 12 and 24 hours to capture the slope of the exponential phase. The metabolite value was calculated as the difference between the metabolite concentration under the exposure condition and the negative control.

Slopes were estimated between 12 and 24 hours for each condition using linear regression, and T-tests were used to compare the slope of the exposure condition with the positive control. To evaluate whether the SCFA has increased proportionally to bacterial growth, pairwise comparisons were conducted using the slope of metabolite levels and growth rates for each exposure condition. Separate two-sample tests were conducted for each SCFA to highlight significant deviations from the positive control.

## 3. RESULTS

### 3.1. *A. muciniphila* MIC for Zn and Co

For Zn, *A. muciniphila* that had been growing in culture for 24 hours (during the exponential growth phase) was inoculated into the modified BHI broth with Zn at 5, 10, 25, 50, 75, 100, 125, 150, 200, 300, 400, 500, 750, 1,000, 1,250, and 1,500 mg/L. Based on OD readings and confirmed by eye, *A. muciniphila* grew at concentrations up to but not including 750mg/L. Because we were interested in concentrations that may affect growth and SCFA production but not stop them, the low concentration was chosen to be 200mg/L, and the high concentration was chosen as 500mg/L, based on our MIC results. For Co, the same procedure was used; however, *A. muciniphila* continued to grow past the maximum dilution of 1,500 mg/L. Thus, we also measured growth at 1,750, 2,000, 2,500, 3,000, 3,500, 4,000, 4,500, and 5,000 mg/L. *A. muciniphila* grew at concentrations up to but not including 4,000mg/L. Based on those findings, the low concentration for Co was chosen to be 200mg/L and the high concentration was chosen to be 3,500mg/L.

### 3.2. *A. muciniphila* Growth with Exposure to Zn and Co

We tested the growth (difference in OD reading between the test and its negative control, denoted with the test condition abbreviation) under each exposure condition in comparison to the positive control at each timepoint. We found that growth was significantly lower under Co+ at 24 (q=0.001) and 48 hours (q=0.001), Co-at 24 hours (q=0.001), and Zn-/Co+ at 24 hours (q=0.035), Zn+/Co+ at 24 hours (q=0.027; **Figure 1**, STable 1). Indicating, that any condition with Co+ reduced growth at the peak of the exponential phase, and the Co-without Zn also reduced growth at the peak of the exponential phase, with growth rebounding after initial declines. All other growth conditions were not significantly different from the positive control, indicating that those exposures did not significantly alter the growth of *A. muciniphila*.

**Figure 1:**
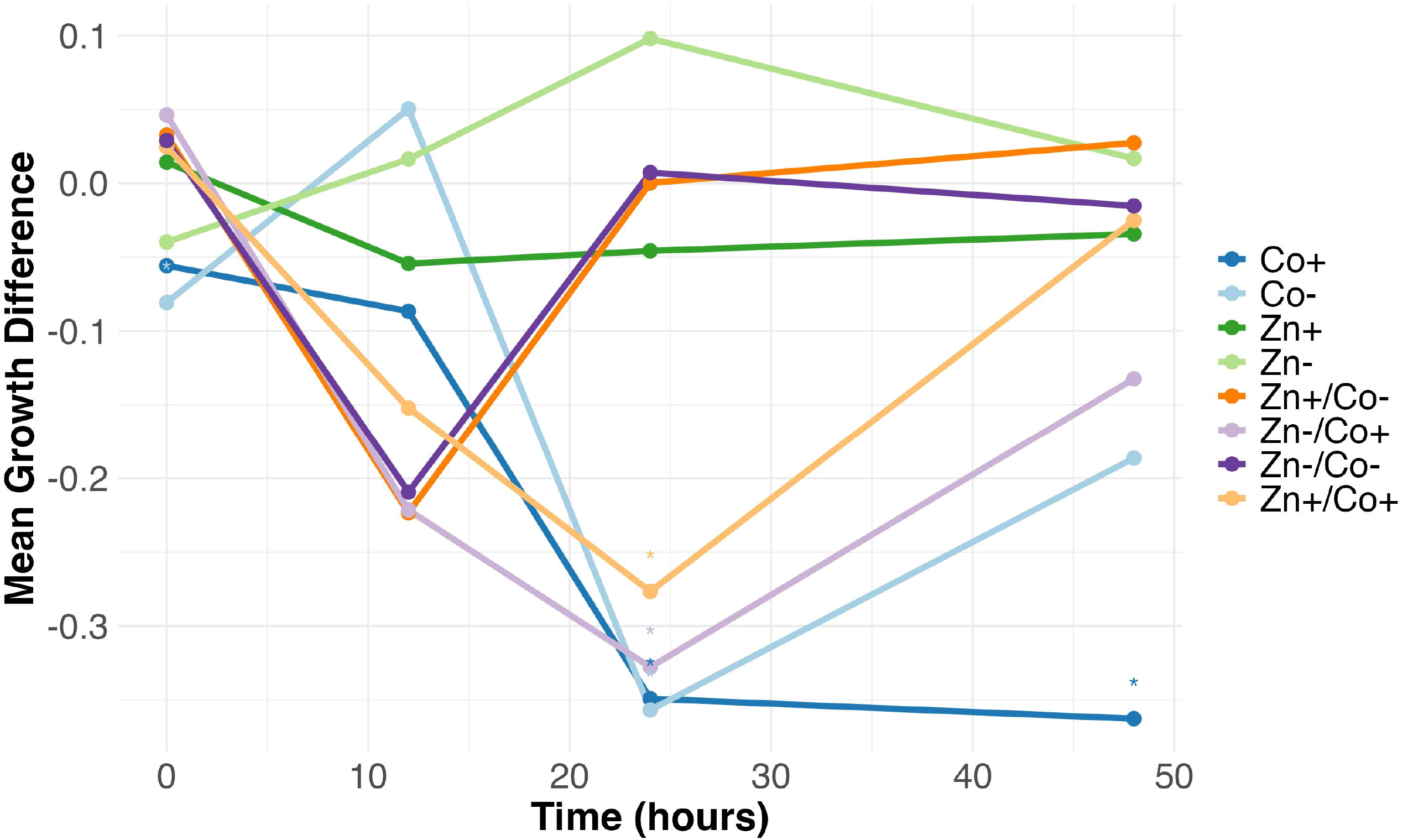
Growth of *A. muciniphila* under each Zn and Co exposure condition from 0-48 hours. The mean growth difference was calculated by subtracting the control values from the exposure values. Zn-is 200mg/L; Zn+ is 500mg/L; Co-is 200mg/L; Co+ is 3,500mg/L. Labels with two metals included both metals at their respective concentrations indicated by the + or – symbols.

### 3.3. *A. muciniphila* SCFA Production with Exposure to Zn and Co

We tested the production rates of propionate, acetate, and butyrate by measuring the difference in slope (denoted with the test condition abbreviation) between each test condition and the positive control over 12 to 24 hours. The primary SCFA produced under the growth conditions used was propionate (**Figure 2**), with significantly lower production of acetate and minimal to no production of butyrate (SFigures 1 & 2). We found that the production rate of propionate was trending upward for Co- (q=0.090), and significantly increased for Zn- (q=0.004), Zn+ (q=0.001), Zn-/Co- (q=0.003), Zn+/Co- (q=0.001), and significantly decreased for Co+ (q<0.001), Zn-/Co+ (q<0.001), and Zn+/Co+ (q<0.001) (Stables 1, 2, 3). The reduced rate of propionate production under Co+ conditions was likely due to the reduced growth observed under these conditions. However, all other exposure conditions had significantly increased rates of propionate production without corresponding increases in growth. We saw no significant differences in the rate of acetate or butyrate production.

**Figure 2:**
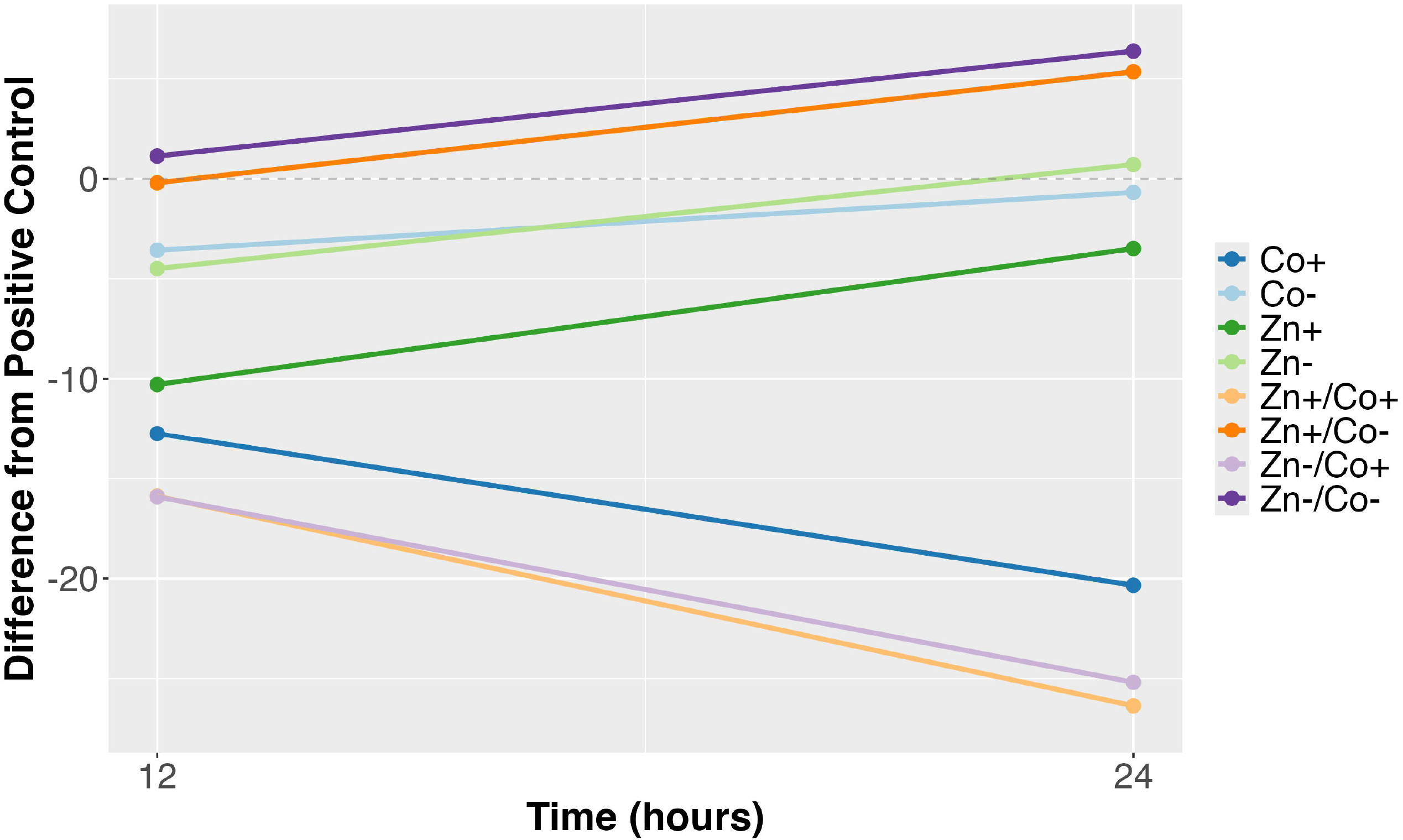
Propionate growth slopes from 12 to 24 hours. Each slope was calculated by subtracting the positive control from the condition’s value. Zn-is 200mg/L; Zn+ is 500mg/L; Co-is 200mg/L; Co+ is 3,500mg/L. Labels with two metals included both metals at their respective concentrations indicated by the + or – symbols.

## 4. DISCUSSION

In this study, we examined the growth and SCFA production of *A. muciniphila* under several Zn and Co exposure conditions. Our MIC results suggest that *A. muciniphila* is highly tolerant to both metals, but especially cobalt, which required much higher concentrations to inhibit growth. Our growth curve analysis supported this finding, with only extremely high concentrations of Co reducing growth. Analysis of SCFAs, primarily propionate, showed increased rates of production under all metal exposure conditions, except very high Co. Considering these results in the context of our epidemiologic study, that indicated a modifying effect of *A. muciniphila* on the association between a prenatal metal signature (high Zn, and low Co, and Cr) and depression in childhood, *A. muciniphila* may be a particularly strong probiotic intervention in this subgroup, given that Zn+ and Co-did not hinder growth, and supported production of SCFAs.

Zn and Co are both essential metals that influence many aspects of human health including neurological function. However, the balance of sufficient yet not excessive exposure is critical to healthy function. (Catalani et al., 2012; Frederickson et al., 2000; Obied et al., 2024; Pfeiffer and Braverman, 1982) Small amounts of exposure can support neurological function, mood regulation, and the health of the gut-brain axis. Primary mechanisms of neurological benefit include Co-B12 synthesis; Co is centrally located in the corrin ring of B12, in which methyl-specific enzymes support metabolic pathways related to myelin formation, DNA methylation, and neurotransmitter regulation. Inhibition of these processes introduces cognitive dysfunction and nerve conduction malfunction (Catalani et al., 2012, Yamada, 2013). Likewise, Zn provides micronutrients that are essential for human life. In the bloodstream, Zn is bound with proteins that interact with plasma-membrane transporters to facilitate passage across the blood-brain barrier. This pathway is essential in antioxidant processes, nerve signaling, modulating synaptic transmission, and regulating neurotransmitters.(Sabouri et al., 2024) Our study found that moderate Zn and Co exposure enhanced SCFA production, which may further support host Zn regulation by increasing stability and structure of proteins that re-distribute Zn in human cells.(Maret, 2009; Sabouri et al., 2024) Together, these findings suggest that appropriate levels of Zn and Co can support neurological function by promoting the growth of *A. muciniphila*, ultimately influencing the gut-brain axis.

While this study adds new information about *A. muciniphila*’s tolerance to Zn and Co, there are some limiting factors to consider. The in vitro nature of this experiment limits our ability to capture the complexities of exposure timing, and biological interactions with the host and other bacteria in the gut microbiome. Thus, the translational value of these results is limited. Future studies should be conducted in in vivo models to better understand the biological interactions involved in toxicity. This study was also limited to the examination of only Zn and Co exposure, and did not include other metals that likely influence the growth and function of *A. muciniphila*. In future studies, additional other metals that humans are regularly exposed to should be considered. Our exploration of metabolic function was also targeted to SCFA production only, not considering other metabolic products of *A. muciniphila* that may influence mental health outcomes in people. Furthermore, the primary SCFA produced in this experiment was propionate, likely due to the ingredients of the enrichment media. Although we gained a sense of altered SCFA production, future studies should also aim to target production of other SCFAs that may have more biological relevance to the translational etiological mechanisms. Finally, future interventional studies should be conducted to evaluate the utility of *A. muciniphila* as an intervention for depression symptoms in sub populations with adverse metal exposures.

## AUTHOR CONTRIBUTIONS

SE contributed to conceptualization, data curation, funding acquisition, methodology, project administration, supervision, and writing – original draft. JE contributed to conceptualization, data curation, formal analysis, funding acquisition, methodology, project administration, and writing – review and editing. MB contributed to formal analysis and writing – original draft. CG contributed to conceptualization and writing – review and editing. VM contributed to conceptualization, formal analysis, methodology, and writing – review and editing.

## FUNDING SOURCES

This work and investigator time was supported by the University of Iowa College of Public Health (New Faculty Research Award), and the National Institute of Environmental Health Sciences (R00ES032884, P30ES005605, P30ES023515). These funders had no role in study design, collection, analysis or interpretation of data, writing of the report or decision to submit the article for publication.

## DATA STATEMENT

All data and code for this analysis are available through Github with Zenodo DOI assignents (made public and listed here upon acceptance).

## Notes

### Competing Interest Statement

The authors have declared no competing interest.

