## Supplemental Material for "Zinc and Cobalt exposure influence *Akkermansia muciniphila* growth and short-chain fatty acid metabolism"

**STable 1:** Growth of *A. muciniphila* under each Zn and Co exposure condition from 0-48 hours.

| Condition | Time | Mean Difference (Condition - Control) | p-value | FDR Corrected q-value |
|---|---|---|---|---|
| ***Co+*** | 0 | -0.056 | 0.264 | 0.621 |
| | 12 | -0.087 | 0.431 | 0.718 |
| | 24 | -0.349 | <0.001 | 0.001 |
| | 48 | -0.363 | <0.001 | <0.001 |
| ***Co-*** | 0 | -0.081 | 0.001 | 0.004 |
| | 12 | 0.050 | 0.630 | 0.775 |
| | 24 | -0.357 | <0.001 | 0.001 |
| | 48 | -0.186 | 0.012 | 0.054 |
| ***Zn+*** | 0 | 0.012 | 0.494 | 0.718 |
| | 12 | -0.054 | 0.588 | 0.752 |
| | 24 | -0.046 | 0.443 | 0.718 |
| | 48 | -0.034 | 0.458 | 0.718 |
| ***Zn-*** | 0 | -0.040 | 0.272 | 0.621 |
| | 12 | 0.016 | 0.871 | 0.929 |
| | 24 | 0.098 | 0.085 | 0.272 |
| | 48 | 0.017 | 0.704 | 0.834 |
| ***Zn+/Co+*** | 0 | 0.025 | 0.543 | 0.752 |
| | 12 | -0.152 | 0.586 | 0.752 |
| | 24 | -0.277 | 0.004 | 0.027 |
| | 48 | -0.025 | 0.787 | 0.869 |
| ***Zn+/Co-*** | 0 | 0.033 | 0.019 | 0.075 |
| | 12 | -0.223 | 0.465 | 0.718 |
| | 24 | <0.001 | 0.997 | 0.997 |
| | 48 | 0.027 | 0.483 | 0.718 |
| ***Zn-/Co+*** | 0 | 0.046 | 0.114 | 0.331 |
| | 12 | -0.222 | 0.323 | 0.689 |
| | 24 | -0.328 | 0.006 | 0.035 |
| | 48 | -0.132 | 0.251 | 0.621 |
| ***Zn-/Co-*** | 0 | 0.029 | 0.050 | 0.176 |
| | 12 | -0.209 | 0.477 | 0.718 |
| | 24 | 0.007 | 0.907 | 0.936 |
| | 48 | -0.015 | 0.748 | 0.855 |

Zn- is 200mg/L; Zn+ is 500mg/L; Co- is 200mg/L; Co+ is 3,500mg/L. Labels with two metals included both metals at their respective concentrations indicated by the + or – symbols.

**STable 2:** Estimate of slope for propionate production by *A. muciniphila* under each Zn and Co exposure condition between 12 and 24 hours. Difference shown is between the slope for that condition and the positive control (with bacteria growth but no metal exposure).

| *Condition* | Estimate | Difference | SE Difference | Z-Statistic | p-value | FDR Corrected q-value |
|---|---|---|---|---|---|---|
| *Co+* | 0.653 | -0.633 | 0.14 | -4.507 | <0.001 | <0.001 |
| *Co-* | 1.526 | 0.241 | 0.142 | 1.697 | 0.09 | 0.09 |
| *Zn-* | 1.719 | 0.433 | 0.149 | 2.91 | 0.004 | 0.004 |
| *Zn+* | 1.852 | 0.566 | 0.155 | 3.643 | <0.001 | 0.001 |
| *Zn-/Co-* | 1.723 | 0.437 | 0.143 | 3.058 | 0.002 | 0.003 |
| *Zn-/Co+* | 0.513 | -0.772 | 0.132 | -5.835 | <0.001 | <0.001 |
| *Zn+/Co-* | 1.748 | 0.463 | 0.133 | 3.476 | 0.001 | 0.001 |
| *Zn+/Co+* | 0.413 | -0.873 | 0.142 | -6.128 | <0.001 | <0.001 |

Zn- is 200mg/L; Zn+ is 500mg/L; Co- is 200mg/L; Co+ is 3,500mg/L. Labels with two metals included both metals at their respective concentrations indicated by the + or – symbols.

**STable 3:** Estimate of slope for acetate production by *A. muciniphila* under each Zn and Co exposure condition between 12 and 24 hours. Difference shown is between the slope for that condition and the positive control (with bacteria growth but no metal exposure).

| *Condition* | Estimate | Difference | SE Difference | Z-Statistic | p-value | FDR Corrected q-value |
|---|---|---|---|---|---|---|
| *Co-* | 0.135 | 0.059 | 0.035 | 1.716 | 0.086 | 0.115 |
| *Co+* | 0.014 | -0.062 | 0.025 | -2.454 | 0.014 | 0.037 |
| *Zn-* | 0.143 | 0.067 | 0.027 | 2.505 | 0.012 | 0.037 |
| *Zn+* | 0.127 | 0.051 | 0.026 | 1.973 | 0.049 | 0.078 |
| *Zn-/Co-* | 0.111 | 0.035 | 0.026 | 1.374 | 0.169 | 0.169 |
| *Zn-/Co+* | 0.016 | -0.06 | 0.025 | -2.357 | 0.018 | 0.037 |
| *Zn+/Co-* | 0.116 | 0.04 | 0.027 | 1.484 | 0.138 | 0.158 |
| *Zn+/Co+* | 0.004 | -0.072 | 0.026 | -2.799 | 0.005 | 0.037 |

Zn- is 200mg/L; Zn+ is 500mg/L; Co- is 200mg/L; Co+ is 3,500mg/L. Labels with two metals included both metals at their respective concentrations indicated by the + or – symbols.

**STable 4:** Estimate of slope for butyrate production by *A. muciniphila* under each Zn and Co exposure condition between 12 and 24 hours. Difference shown is between the slope for that condition and the positive control (with bacteria growth but no metal exposure).

| *Condition* | Estimate | Difference | SE Difference | Z-Statistic | p-value | FDR Corrected q-value |
|---|---|---|---|---|---|---|
| *Co-* | 0.006 | 0.01 | 0.002 | 5.258 | <0.001 | <0.001 |
| *Co+* | -0.016 | -0.013 | 0.001 | -8.851 | <0.001 | <0.001 |
| *Zn-* | 0.007 | 0.011 | 0.003 | 3.913 | <0.001 | <0.001 |
| *Zn+* | 0.008 | 0.012 | 0.002 | 5.505 | <0.001 | <0.001 |
| *Zn-/Co-* | 0.004 | 0.008 | 0.002 | 3.393 | 0.001 | 0.001 |
| *Zn-/Co+* | -0.014 | -0.01 | 0.003 | -3.199 | 0.001 | 0.001 |
| *Zn+/Co-* | 0.012 | 0.015 | 0.001 | 10.904 | <0.001 | <0.001 |
| *Zn+/Co+* | -0.019 | -0.015 | 0.004 | -3.582 | <0.001 | <0.001 |

Co- is 200mg/L; Co+ is 3,500mg/L; Zn- is 200mg/L; Zn+ is 500mg/L. Labels with two metals included both metals at their respective concentrations indicated by the + or – symbols.

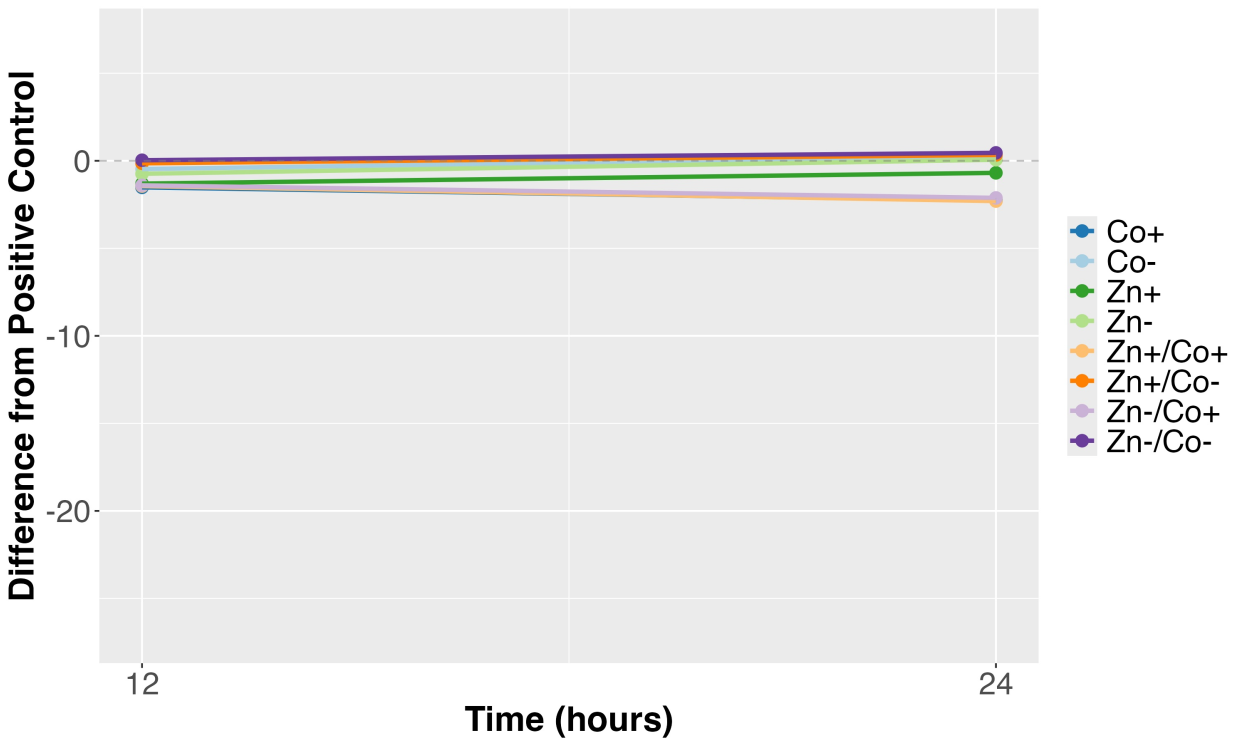


**SFigure 1:** Acetate growth slopes from 12 to 24 hours. Each slope was calculated using the positive control subtracted from the condition's value. Co- is 200mg/L; Co+ is 3,500mg/L; Zn- is 200mg/L; Zn+ is 500mg/L. Labels with two metals included both metals at their respective concentrations indicated by the + or – symbols.

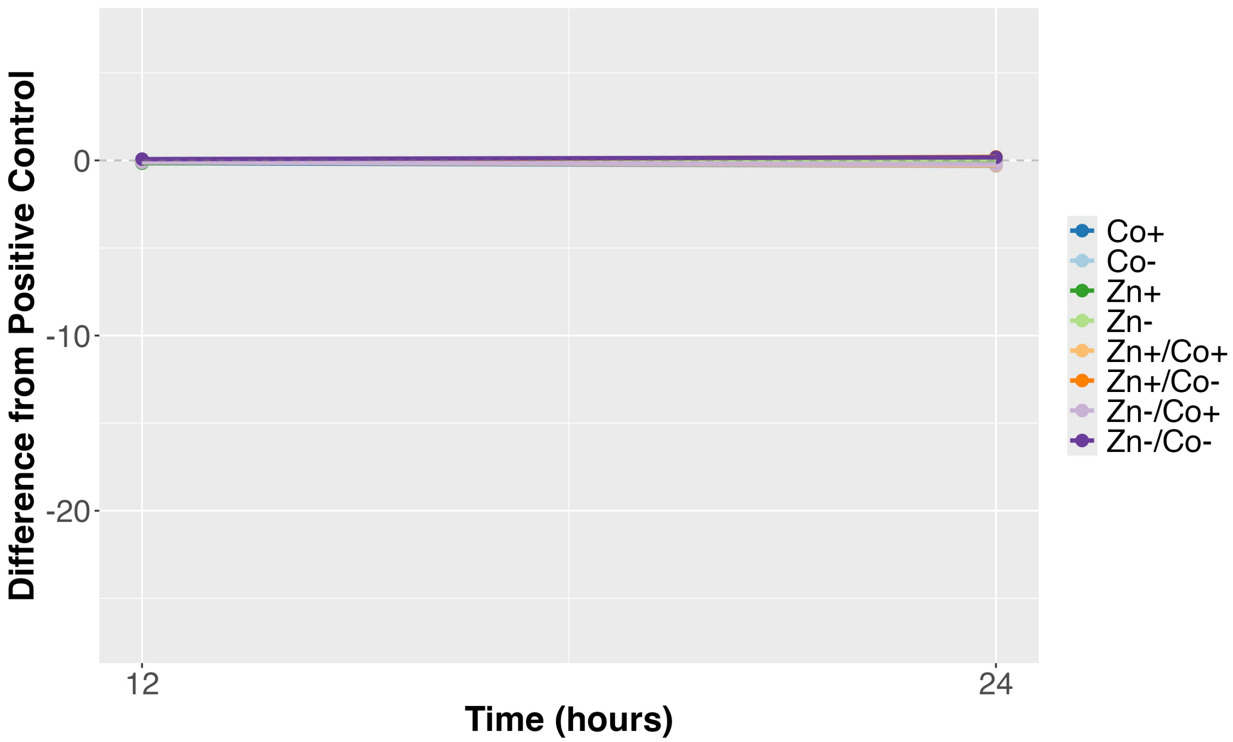


**SFigure 2:** Acetate growth slopes from 12 to 24 hours. Each slope was calculated using the positive control subtracted from the condition's value. Co- is 200mg/L; Co+ is 3,500mg/L; Zn- is 200mg/L; Zn+ is 500mg/L. Labels with two metals included both metals at their respective concentrations indicated by the + or – symbols.
